# Pliable synaptic fidelity and excitatory inter-synaptic crosstalk in the intact brain

**DOI:** 10.1101/2022.03.04.483016

**Authors:** James P. Reynolds, Thomas P. Jensen, Sylvain Rama, Kaiyu Zheng, Leonid P. Savtchenko, Dmitri A. Rusakov

**Affiliations:** UCL Queen Square Institute of Neurology, University College London, United Kingdom

## Abstract

Memory formation in neural circuits may involve changes in synaptic efficacy and in cell intrinsic excitability, yet how this process unfolds in the living brain has remained elusive. Here, we employed multiplexed imaging of genetically encoded indicators of glutamate and Ca^2+^ in mouse barrel cortex to detect increased fidelity coupled with reduced excitation of thalamocortical connections that undergo whisker-stimulation induced LTP. High-resolution imaging revealed that whisker stimuli trigger excitatory synaptic activity that generates extrasynaptic glutamate transients reaching the bulk of neighbouring synapses in the target cortical area. Our findings pave the way to understanding basic plasticity features of the synaptic connectome while revealing a significant component of volume-transmitted glutamatergic signalling among cells in the intact brain.

**One Sentence Summary:** Sensory-stimulation LTP increases fidelity while reducing excitation at individual thalamocortical connections which generate spatially intersecting glutamate discharges

---

Excitatory glutamatergic synapses underpin the workings of the brain. Their ability to change transmission strength in a coincidence-dependent manner has been considered key to the process of learning described by Hebb (*1*). Hebbian principles have found their dependable empirical prototype in the synaptic long-term potentiation (LTP) (*2*), which has since been associated with memory trace formation (*3-7*). However, how the build-up of synaptic potentiation evades runaway network excitation - either through homeostatic synaptic weight re-scaling (*8-10*), or through altered cell excitability (*11-13*), or both (*14-16*) - has been disputed. The issue remains debatable because separating out synaptic efficacy changes from the effects of cell and network excitability in vivo has been difficult. In practice, LTP in the intact brain has been shown almost exclusively in the bulk of synapses, which limits our understanding of whether it involves an increase in the postsynaptic receptor current or in release probability (P_r_), or both.

In general, P_r_ of small excitatory synapses can vary >5-fold along individual axons (*17*) or dendritic trees (*18*), and is highly use-dependent (*19*), whereas its average value in vivo is generally low (*20*). These features are in line with a theory that relates optimal information handling in the cortex predominantly to P_r_ control (*21, 22*). In brain slices, direct P_r_ readout can be achieved by imaging either postsynaptic Ca^2+^ transients (*23, 24*) or presynaptic glutamate release (*17, 25*), but such methods rely on full control of action potentials, which could be unattainable for highly interconnected in vivo networks.

We sought to address some of these long-standing issues, firstly, by establishing a P_r_ tracking method that would not require spiking control in vivo, and secondly, by examining transmission at individual thalamocortical (TC) synapses in the course of LTP induced by contralateral rhythmic whisker stimulation (RWS-LTP) (*26-28*). To test and validate P_r_ tracking, we took advantage of a previously established multiplexed imaging technique (*17*). In organotypic hippocampal slices, we patched a CA3 pyramidal cell expressing the optical glutamate sensor iGluSnFR (Fig. 1A, fig. S1A; Methods). After whole cell break-in and dialysis of the cell with the red-shifted Ca^2+^ indicator Cal-590 (in some cases also with the tracer Alexa Fluor 594), we focused on individual presynaptic boutons traced along the axon, for high-speed two-photon excitation (2PE) ‘Tornado scan’ imaging (Fig. 1B, fig. S1B; Methods). Four action potentials evoked at the soma produced clear multiplexed readout of evoked glutamate release and presynaptic Ca^2+^ entry (Fig. 1C; fig. S1C). The corresponding dynamics of absolute Ca^2+^ concentration ([Ca^2+^]) was assessed using Ca^2+^ sensitivity of the Cal-590 fluorescence lifetime (Fig. 1D), as detailed previously (*17, 29*).

**Figure 1.**
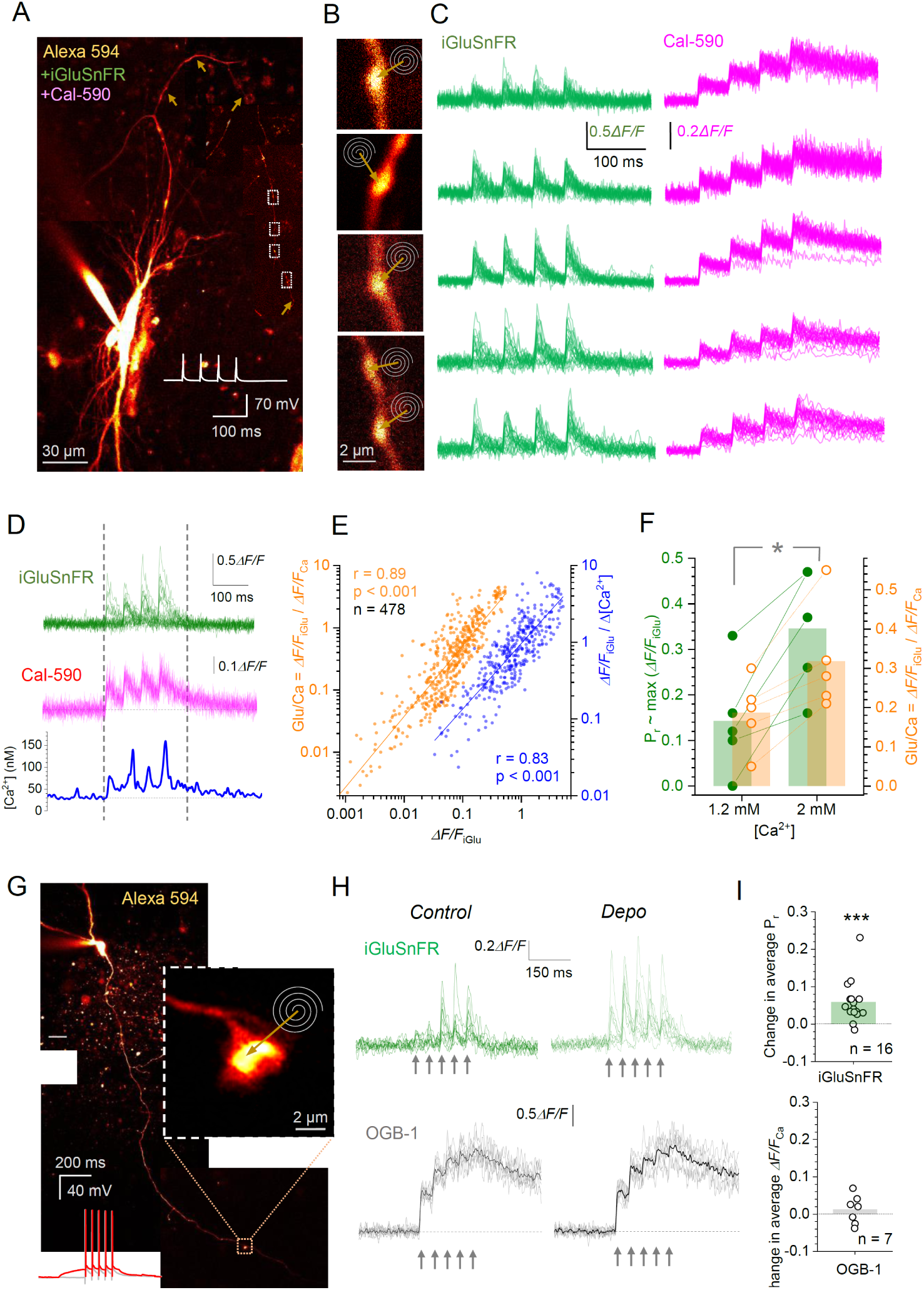
The ratio of glutamate versus Ca^2+^ fluorescence signals at spiking presynaptic boutons can faithfully track changes in synaptic release probability P_r_. (**A**) CA3 pyramidal cell (organotypic hippocampal slice) expressing SF-iGluSnFR.A184V and dialysed whole-cell with Cal-590 and 4 μM Alexa-594, with 4 ROIs along the axon (arrowheads, dotted rectangles; collage of 8-17 μm deep image stack projections, λ_x_^2P^=800 nm for tracing; example with iGluSnFR tracing in fig, S1A-B); inset trace, four action potentials 50 ms apart elicited at the soma in current-clamp. (**B**) Presynaptic axonal boutons (ROIs shown in A); spirals and arrows, Tornado scan positions. (**C**) Fluorescence time course (integrated Tornado scan signal, 500hz sampling (single boutons) or 333Hz (multiple boutons) in iGluSnFR (green) and Cal-590 (magenta) channels (λ_x_^2P^=910 nm), in response to four action potentials 50 ms apart, as indicated, for the boutons in B; 20-24 trials shown. (**D**) Example of fluorescence responses as in C (green and magenta), with the FLIM-based readout of presynaptic [Ca^2+^] (blue; bottom panel); vertical lines, sampling window for *ΔF/F*_*0*_ value averaging. (**E**) The ratio of *ΔF/F* iGluSnFR response (250 ms over four responses, as in D) versus either *ΔF/F* Cal-590 response (‘Glu/Ca’ ratio, orange) or FLIM readout of [Ca^2+^] (blue), plotted against *ΔF/F* iGluSnFR response (direct glutamate release readout, n = 478 boutons); straight lines, linear regression; r, coefficient of correlation; log-log scale. (**F)** *ΔF/F* iGluSnFR (green, scales with P_r_) and Glu/Ca responses (orange) at two concentrations of extracellular Ca^2+^, as indicated; dots, individual experiments; bars, average values; *, p < 0.01 (n = 5), in both cases. (**G**) Fragment of dentate gyrus granule cell dialysed whole-cell with OGB-1, with a tracked giant mossy fibre bouton (dotted rectangle; collage of 10-15 μm deep image stack projections) and Tornado scan position indicated (inset); trace, five action potentials elicited at the soma in current-clamp, at V_m_= -70 mV (grey) and during 200 ms depolarisation by ∼15 mV (red). (**H**) Example of fluorescence responses in mossy fibre boutons expressing iGluSnFR (top, 11 trials shown) and those filled with OGB-1 (bottom, 10 trials; black line, average) during five action potentials (arrows), in control conditions and under somatic depolarisation, as shown by trace in G. (**I**) Changes in P_r_, as gauged by the failure rate of evoked iGluSnFR signals (top, n = 16 boutons), and changes in presynaptic Ca^2+^ entry (*ΔF/F* OGB-1, bottom; n = 7 boutons); dots, individual experiments; bars, average; ***, p < 0.001.

In these experiments, iGluSnFR signal varied many-fold, reflecting variable quantal content (including failures) of release events (Fig. 1A-D); this variability was not due to recording instability because it was several times smaller in the matching glutamate 2PE-uncaging tests (fig. S1D-F). At the same time, the spike-evoked Ca^2+^ signal was remarkably stable, suggesting that, in conditions of non-saturation, it would scale with the number of action potentials, a notion consistent with somatic Ca^2+^ imaging in vivo (*30*). Thus, for a given synapse, the ratio between glutamate- and Ca^2+^-sensitive signals (Glu/Ca) should scale with the amount of glutamate released per action potential, or synaptic release efficacy proportional to P_r_. Indeed, when we plotted the Glu/Ca ratio (or the ratio Glu / [Ca^2+^] entry) against direct P_r_ readout (iGluSnFR signal amplitude), the relationship showed excellent linear regression (Fig. 1E).

To test if the Glu/Ca ratio would follow P_r_ changes induced in the same synapse, we examined two cases, one with Ca^2+^-dependent and one with Ca^2+^-independent modulation of P_r_. In the first case, we changed extracellular [Ca^2+^] between 1.2 and 2.0 mM and documented the corresponding change in P_r_ at CA3-CA1 synapses using iGluSnFR readout (Fig. 1F, left ordinate). Here, Glu/Ca reliably reported P_r_ increases in all tested synapses (Fig. 1F, right ordinate), reflecting the highly supralinear dependence of P_r_ on Ca^2+^ entry hence extracellular [Ca^2+^] (*31*). We also confirmed that at 2.0 mM extracellular [Ca^2+^], Glu/Ca scaled linearly with iGluSnFR readout (fig. SG). In the second case, we monitored Ca^2+^ transients and glutamate release from giant mossy fibre boutons tracked from dentate granule cell somata (Fig. 1G). In this circuitry, presynaptic depolarisation can increase glutamate release (*32*), which appears dissociated from changes in presynaptic Ca^2+^ entry (*33*). Indeed, depolarising the soma by ∼15 mV boosted glutamate release in response to five action potentials (Fig. 1H) whereas Ca^2+^ entry remained unchanged (Fig. 1I), at various distances from the soma (fig. S1H). These tests confirmed that, at least for relatively short spike bursts, the Glu/Ca ratio would follow the amount of glutamate released per spike, thus providing P_r_ readout when spiking cannot be directly monitored.

Equipped with this approach, we sought to monitor P_r_ changes in TC projections that respond to contralateral rhythmic whisker stimulation (RWS) in vivo. We targeted the circuitry of the posterior medial nucleus (POm) of the thalamus, monitoring axonal projections to the barrel cortex in S1 (*34-36*), where ramifications are widespread but sparse (*37, 38*). To enable the multiplexed imaging of presynaptic Ca^2+^ and glutamate release at individual connections, we virally transduced two genetically-encoded indicators, red-shifted cytosolic jRGECO1a (*39*) for Ca^2+^ and green surface-bound iGluSnFR for glutamate, in POm neurons (synapsin promoter) and S1BF astroglia (GFAP promoter), respectively (Fig. 2A), using a method established previously (*36, 40*) (Methods). Two weeks after AAV transduction, a cranial window was implanted over S1BF, so that individual presynaptic boutons of the jRGECO1a-lablled TC axons could be traced once the animal has recovered (Fig. 2A).

**Figure 2.**
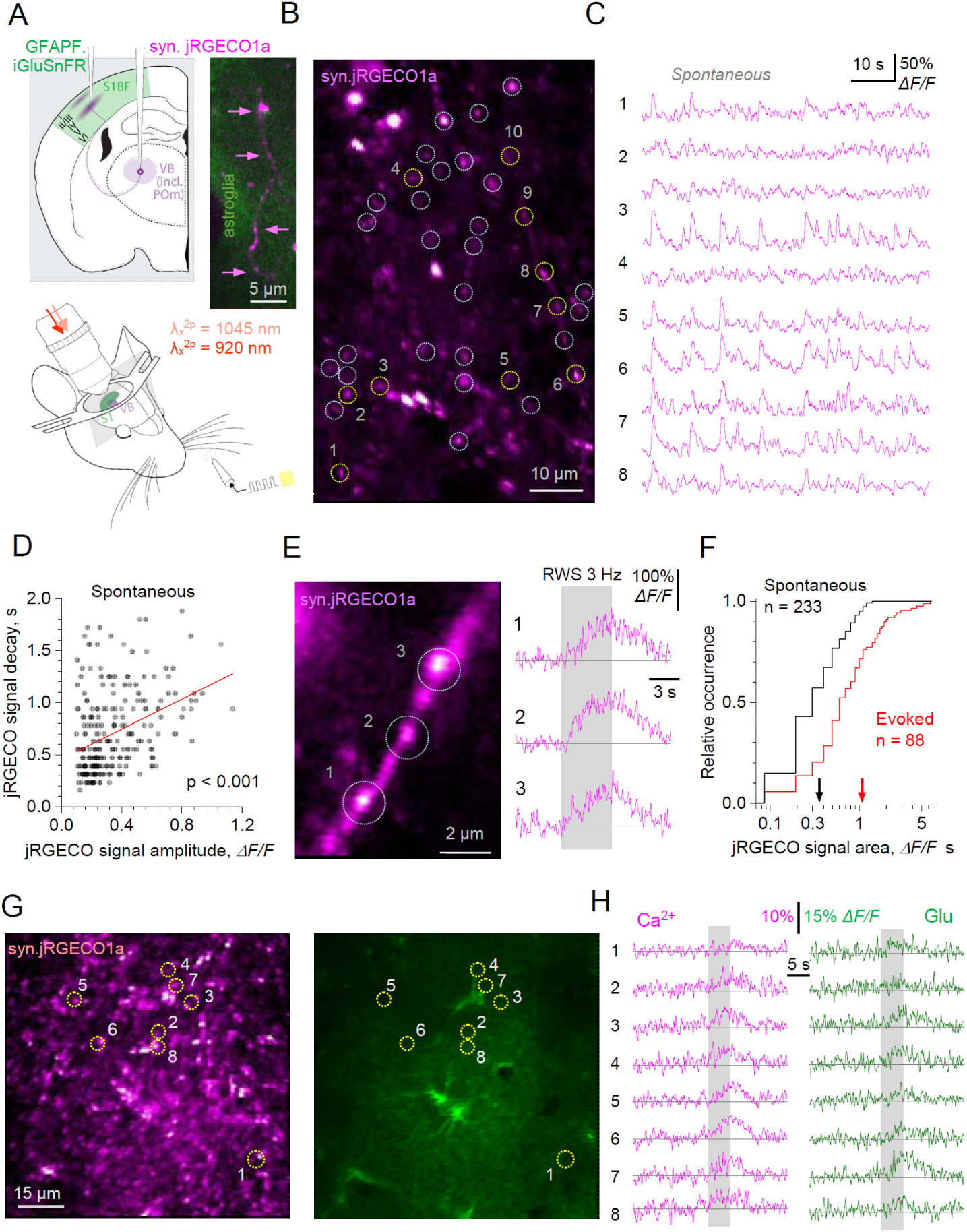
Presynaptic Ca^2+^ imaging in thalamocortical (TC) axonal boutons covers the dynamic range of axonal firing enabling Pr tracking in vivo using the Glu/Ca ratio. (**A**) Experimental arrangement illustrating viral transduction of jRGECO1a and iGluSnFR.A184S in the barrel cortex S1BF astroglia, and thalamus VB neurons, respectively (top); with a cranial window, under dual length two-photon excitation regime, as illustrated (bottom); inset image, snapshot through the cranial window depicting expression of jRGECO1a in TC axons (magenta, arrowheads) and iGluSnFR in local astroglia (green); see Methods. (**B**) A wider-frame snapshot (jRGECO1a channel; depth ∼100 μm, layer 2-3 S1BF) showing tentative TC axonal boutons (selected arbitrarily, based on morphology; dotted circles). (**C**) Snapshot of spontaneous Ca^2+^ activity recorded in numbered boutons shown in B (other candidate boutons appeared silent). (**D**) The decay of spontaneous presynaptic Ca^2+^ signals recorded as in B-C, plotted against the *ΔF/F*_0_ signal amplitude (jRGECO1a fluorescence, n = 233 events); line, linear regression (significant slope at p < 0.001). (**E**) An example of TC axonal fragment with three jRGECO1a-expressing boutons (1-3, left) and their fluorescence responses to a contralateral rhythmic whisker stimulation (RWS, 5 s at 3 Hz; grey segment, right). (**F**) Frequency distribution of the presynaptic signal strength (area under the ΔF/F0 curve) for spontaneous and evoked events at TC boutons, as indicated. (**G**) Example of multiplexed-imaging snapshots in the jRGECO1a (left) and iGluSnFR (right) channels, with tentative TC axonal boutons (dotted circular ROIs) selected based on the detectable presynaptic Ca^2+^ and glutamate release activity. (**H**) Fluorescence responses in TC axonal boutons shown in G (area-integrated in ROIs 1-8 in G) to contralateral RWS (5 s at 3 Hz, grey area), as indicated.

First, we explored the dynamic range of presynaptic Ca^2+^ activity in vivo, to probe saturation of jREGO1a readout. Awake animals showed spontaneous Ca^2+^ activity in 20-30% tentative axonal boutons that we traced through the cranial window using cytosolic jREGO1a fluorescence (Fig. 2B-C). At individual boutons, the signal amplitude was correlated with its decay (Fig. 2D; analysis detail in fig. S2A), with signal strength increasing with its frequency (fig. S2B). The latter suggested the occurrence of variable-intensity spike bursts rather than uniform events. Elevating the anaesthesia level visibly suppressed spontaneous activity, by ∼85% (Methods), thus enabling us to record, without contamination, synaptic responses to 5 s-long RWS at 3 Hz (Fig. 2E). Overall, the evoked Ca^2+^ responses were 3-4 times stronger than individual spontaneous events, but in either case the average Ca^2+^ signal was 4-5 times smaller than the largest (Fig. 2F). This suggested, firstly, no appreciable jREGO1a signal saturation for RWS lasting 5 s or less, and secondly, that RWS-evoked responses represented multiple-spike bursts.

We therefore switched to multiplexed imaging mode (Fig. 2G), detecting RWS-evoked glutamate release in some (20-30%) of the TC axonal boutons that show Ca^2+^ activity (Fig. 2H). Indeed, multiple-trial recordings showed much lower failure rates for Ca^2+^ compared with glutamate signal (fig. S2C-E), consistent with the stochastic nature of its release and/or detection failure due to the prevalence of glutamate uptake (see below). We next focused on individual boutons using Tornado scan regime and documented RWS-evoked responses in Ca^2+^ and glutamate channels (Fig. 3A-B), in control conditions and 10-30 min following the RWS-LTP induction protocol (5 Hz for 120s; Fig. 2C). In individual boutons, LTP induction was followed by a significant increase in the Glu/Ca ratio (by 47 ± 12%, p < 0.001, n = 19 boutons in 6 animals; Fig. 3D, fig. S3A-B), reflecting the increased average synaptic P_r_. Consistent with optical quantal-analysis studies in slices (*24*), the smallest P_r_ readout was correlated with the biggest change post-LTP (Fig. 3E).

**Figure 3.**
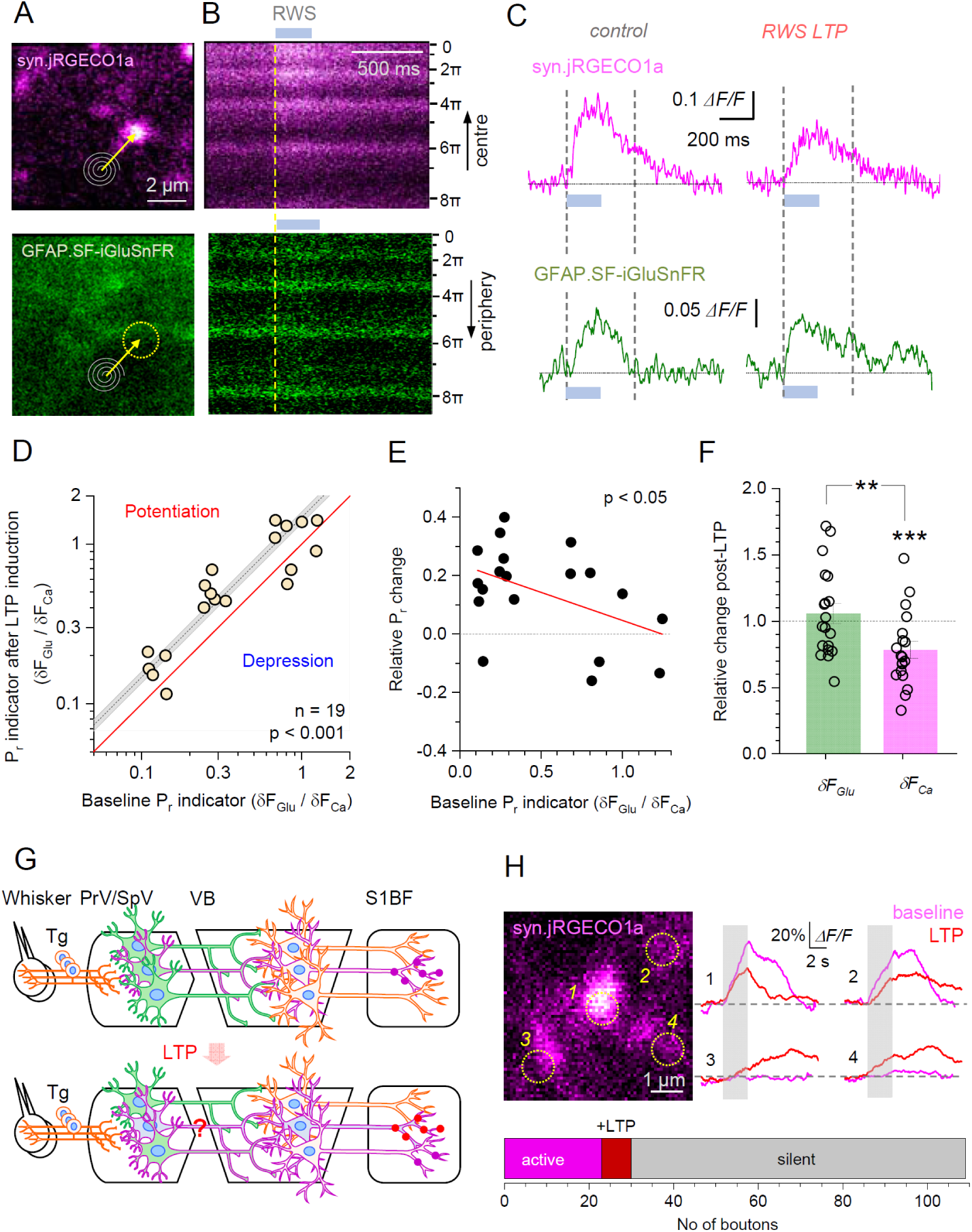
Increased fidelity and decreased excitability of TC synapses following rhythmic-whisker-stimulation induced long-term potentiation (RWS-LTP). (**A**) A snapshot of multiplexed frame imaging, with a jRGECO1a-expressing TC axonal bouton (top, magenta) and local iGluSnFR-expressing astroglia (bottom, green; dotted circle, bouton of interest ROI); spirals and arrows, Tornado scan position. (**B**) Tornado scan example for the bouton in A, showing a fluorescent transient upon 200 ms RWS (grey segment; dotted line, onset) in both channels; ordinate, rotation angle of the Tornado spiral (rad). (**C**) Bouton-average (Tornado-scan integrated, ∼500 Hz sampling) fluorescence time course in jRGECO1a and iGluSnFR channels, as indicated, during 200 ms RWS stimulation (grey segment), before (left) and 10-30 min after RWS-LTP induction protocol (right); average of four trials. Dotted lines, sampling windows to calculate the Glu/Ca ratio (using areas under the *ΔF/F*_0_ curves). (**D**) Statistical summary of experiments shown in A-C: Glu/Ca ratio 10-30 min after LTP induction plotted against Glu/Ca ratio in control condition; solid line, no-change boundary; dots, individual boutons (n = 19 boutons in 6 animals); dotted line and grey segment, mean ± SEM; log-log scale; p < 0.001, H_0_ hypothesis (no change) rejection level, one-sample t-test; further data presentation in fig. S3A-B. (**E**) Relative change in Glu/Ca ratio upon RWS-LTP induction, plotted against baseline Glu/Ca, dataset as in D; solid line, linear regression (slope, p < 0.05). (**F**) Relative changes, upon RWS-LTP, in the bouton-average *ΔF/F*_*0*_ responses recorded in the iGluSnFR (green) and jRGECO1a (magenta) channels, as in C; dots, individual boutons (n = 19), bars, mean ± SEM; **, p < 0.01; ***, p < 0.005. (**G**) Diagram of neural circuitry that could be activated by whisker stimulation, in control conditions (top) and after RWS-LTP induction (bottom). Notations: magenta dots and cell boundaries, activated cells and synapses; PrV/SpV, principal and spinal nuclei of the trigeminal complex (brainstem); POm, posterior medial nucleus of the thalamus); S1BF (barrel cortex layers 2-3). (**H**) Image, example of four arbitrarily selected TC axonal bouton (jRGECO channel); traces: boutons 1 and 2 show robust Ca^2+^ response to RWS (grey segment), which is reduced 10-20 min after RWS-LTP; boutons 3 and 4 show Ca^2+^ response after but not before RWS-LTP induction; graph, summary: among arbitrarily selected tentative TC boutons 96 were Ca^2+^-silent throughout (grey), 23 were active before (magenta) and additional 7 (red) after RWS-LTP induction (data from 3 animals).

Surprisingly, the Glu/Ca ratio increase was mainly due to a decrease in the presynaptic Ca^2+^ signal rather than an increase in glutamate transients (Fig. 3F). This was not because of the differential photobleaching of jRGECO1a versus iGluSnFR: both channels revealed a similarly small (∼12%) decrease in the baseline fluorescence post-induction (fig. S3C). Thus, the data appeared to suggest that RWS-LTP induction lowered excitability of the projecting VB neurons, or possibly of the upstream primary trigeminal nucleus neurons, or both, while providing their unchanged excitatory drive (the amount of glutamate discharged) in the barrel cortex for the same RWS (Fig. 3G). However, RWS-LTP is known to boost postsynaptic responses in cortical neurons (*26, 28*). We therefore hypothesised that RWS-LTP, while reducing excitability of the active pathway, increases excitability of neuronal projections that were silent in control conditions (Fig. 3G). In this case, we should be able to detect an increase in the numbers of Ca^2+^-active axonal boutons after LTP induction. We therefore monitored Ca^2+^ activity in 119 tentative axonal boutons (randomly selected, by their morphology only), and found that 23 were active in control conditions whereas additional 7 became active post-LTP (Fig. 3H). This observation suggests that there could be a redistribution of excitability among cortical networks during RWS-induced plasticity, a phenomenon predicted earlier based on current-source density mapping (*26*).

Indeed, modulation of intrinsic cell excitability has emerged as an important constituent of memory trace formation (*12, 14, 41-44*), and several mechanisms have been reported which underpin activity-dependent bidirectional excitability changes in cortical neurons (*45, 46*). The decreased intensity of evoked synaptic discharges at active TC synapses during LTP found here (Fig. 3F) could reflect the inverse relationship between recent cell firing activity and its excitability, possibly via regulation of the *I*_*h*_ current (*41, 47*) or K^+^ conductance (*48, 49*), as shown in various neuronal types. Conversely, the LTP-related increase in the number of active TC synapses (Fig. 3H) could potentially be explained by the increased excitability of thalamic neurons projecting to the cortex (Fig. 3G) after repetitive subthreshold activation of their glutamate inputs, as found in other settings (*50-52*).

It has recently transpired that, in similar experimental settings, LTP could drive glutamate transporter expressed by local astroglia away from potentiated synapses (*40*). Although this appears to boost extrasynaptic glutamate escape in ex vivo preparations (*40, 53*), little is known about extrasynaptic glutamate escape in the intact brain. The notion of glutamate-mediated synaptic crosstalk among central synapses has been contentious, despite evidence in brain slices (*54-58*), because synaptic connectivity is traditionally considered as ‘wired’ circuitry, akin to that in computer chips.

To address this, we imaged jRGECO1a-filled axonal boutons using 2D reconstructions of Tornado scans (*17*) (Fig. 4A, top) while recording landscapes of iGluSnFR fluorescence in the surrounding astroglia (Fig. 4A, bottom). The iGluSnFR landscapes were averaged over two 200 ms time windows, one before and one during the RWS-induced response (Fig. 4B). The numerical ratio between the two corresponding images thus provided a 2D landscape of *ΔF/F*_*0*_ readout reflecting the extracellular 2D glutamate transient profile, at and near the synapse (Fig. 4C, top). In most cases, the *ΔF/F*_*0*_ image revealed a hotspot, pointing to the tentative glutamate release site, with the signal fading with distance (Fig. 4; fig. S4A). The analysis of these images suggested, firstly, that in control conditions the RWS-evoked glutamate transient extends for at least ∼2.5 μm from the release site, with a biexponential decay (Fig. 4D, left). Secondly, that RWS-LTP induction does not change its decay profile within ∼0.5 μm from the release site while increasing significantly its longer-range component (at >1.5 μm, p < 0.02; Fig. 4D-E).

**Figure 4.**
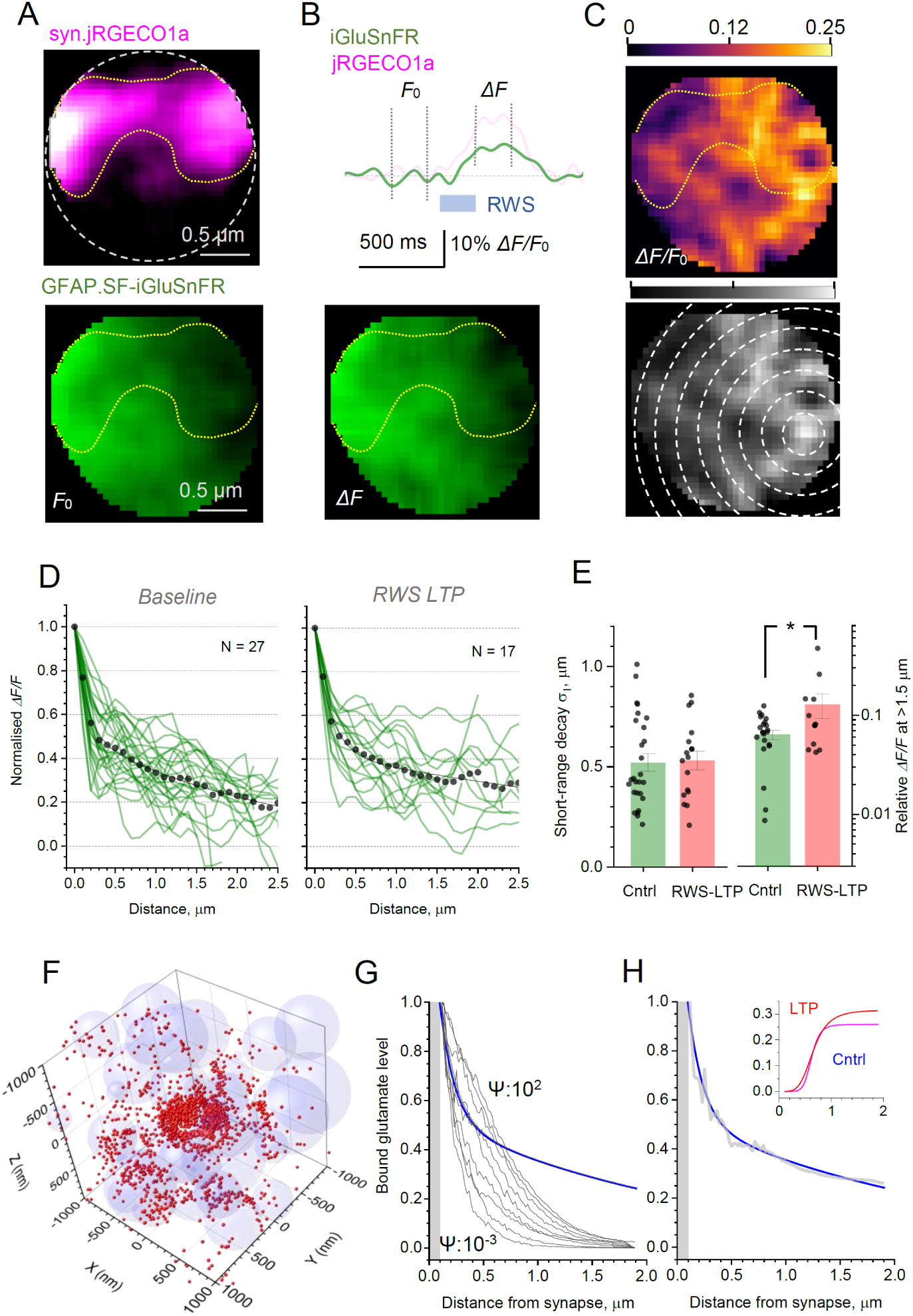
Extrasynaptic glutamate signalling in TC axonal boutons in response to whisker stimulation. **(A)** A snapshot of high-resolution Tornado-scan multiplexed imaging, showing a jRGECO1a-expressing TC axonal bouton (top, magenta) and the surrounding iGluSnFR-expressing astroglia (bottom, green), in resting conditions (200 ms average at 500 Hz sampling); dotted circle, Tornado scan area coverage; yellow dotted line, axonal bouton morphology outline as judged by the fluorescence of cytosolic jRGECO1a. (B) Traces, Tornado scan-integrated fluorescence time course in the iGluSnFR channel (green; jRGECO1a magenta channel shown for illustration only), during 200 ms RWS stimulation (grey segment); vertical dotted lines, sampling windows (resting and signal intervals) for generating the *ΔF/F*_*0*_ iGluSnFR signal landscape. Image (bottom), landscape of iGluSnFR fluorescence as in A, but during the evoked signal (200 ms average at 500 Hz sampling). (C) Top, the *ΔF/F*_*0*_ iGluSnFR signal landscape generated as the ratio of *ΔF* image in B and *F*_0_ Image in A; bar, false colour signal scale; arrow, hotspot (fluorescence peak). Bottom, image as above, but in calibrated grey scale, with concentric circles to calculate average signal decay from the hotspot (further examples in fig. S4A). (D) Normalised decay profile of the *ΔF/F*_*0*_ iGluSnFR signal, with respect to the hotspot, in baseline conditions (left) and 10-30min after RWS-LTP induction (right); green lines, individual axonal boutons; dots, average data; solid line, best-fit biexponential approximation in the form *A*_*1*_·exp(-x/λ_1_) + *A*_2_·exp(-x/λ _2_); control best-fit shown: *A*_1_ = 0.491, λ_1_ = 0.134 μm, *A*_2_ = 0.518, λ_2_ = 2.38 μm; LTP best-fit: *A*_1_ = 0.572, λ_1_ = 0.181 μm, *A*_2_ = 0.431, λ_2_ = 5.42 μm. (E) Short-range decay constant (left) and the average *ΔF/F*_*0*_ iGluSnFR signal value for distances >1.5 μm form the hotspot, compared for control conditions and post-LTP, as indicated; dots, individual bouton data; bars, mean ± SEM; *, p = 0.024. (F) Simulation snapshot (2 μm wide fragment of the 4 μm simulation arena) of release, diffusion, and binding of glutamate molecules (red dots) to astroglia-expressed iGluSnFR (and glutamate transporters GLT-1), in the neuropil modelled *in silico* by a scatter of overlapping spheres (text and Methods); for clarity, only spheroid shapes representing astroglia are shown (see fig. S4C-D for all-component illustrations); free diffusivity *D* = 0.5 μm^2^/ms, glutamate-indicator binding efficiency parameter (Methods) Ψ = 0.1 ms; time point, 4 ms post-release. (G) Black lines, simulated concentration profiles of iGluSnFR-bound glutamate, relative to the release site (grey segment, synaptic cleft area), over the range of Ψ (0.001, 0.01, 0.1, 0.3, 0.5, 0.75, 1, 10, 100 ms); lowest and highest Ψ values shown; blue line, experimental profile of the *ΔF/F*_*0*_ iGluSnFR signal, as in D (control conditions). (H) Experimental profile as in G (blue, main graph), with the best-fit simulated profile (grey) at binding efficiency Ψ = 0.5 ms in the initial part (<0.7 μm), and a best-fit tail addition (at >0.7μm) shown in the inset graph, in control condition (blue) and after RWS-LTP (red).

In this brain region, the volume density of cortical-cortical (CC) and thalamocortical (TC) synapses is, respectively, ∼1.10 and ∼0.23 μm^-3^ (*59, 60*), giving a random-scatter average nearest-neighbour distance (*61*) of 0.90 μm between TC synapses, and 0.53 μm between any synapses. Thus, the average *ΔF/F*_0_ decay profile (Fig. 4D), which reflects glutamate bound to iGluSnFR post-release, points to significant inter-synaptic crosstalk involving glutamate receptors with the affinity comparable with iGluSnFR or higher. To understand whether such crosstalk can be generated by an individual (recorded) synapse or whether it reflects multiple synaptic activation in the local area, we modelled a 4 μm cube of the barrel cortex neuropil, by expanding a previously established Monte Carlo approach (*62, 63*). The tissue was modelled using a random scatter of unequal overlapping spheres (*64*) that would represent cortical astroglia taking ∼10% tissue volume fraction (*65*) and neuronal structures taking ∼70%, leaving free ∼20% volume (space filling *β* = 0.8) for the extracellular space (*66, 67*) (Fig. 4F; Methods).

Glutamate molecules were ‘released’ inside a 220 nm-wide, 20 nm-high synaptic cleft (*59*), diffusing with *D*=0.5 μm^2^/ms measured previously with anisotropy-FLIM (*68*). First, we validated the approach by comparing the outcome of Monte Carlo simulations of freely diffusing molecules with the theoretical value of Maxwell’s diffusivity (0.09 μm^2^/ms for *β* = 0.8) in a similar porous medium (Fig. 4B). Next, we controlled the binding of molecules to the astroglia-representing spheroid shapes using time constant (integrated parameter) Ψ that represents the unknown local concentration of glutamate-binding entities, such as iGluSnFR and the main glial glutamate transporter GLT-1 (*69, 70*) (Fig. 4F, fig. S4C-D; Methods). Thus, Ψ was the only free parameter to fit the simulated distribution of bound particles to the experimental *ΔF/F*_0_ iGluSnFR profile (shown in Fig. 4D). Intriguingly, varying Ψ value 10^6^ -fold (from no binding to 100% instantaneous binding) consistently predicted a shorter glutamate diffusion spread than seen experimentally (Fig. 4G). The consistent mismatch suggested that glutamate signal at >0.5 μm from the synapse should come mainly from neighbouring sources. This missing signal ‘tail’ was well approximated by a Hill function, showing a ∼20% increase after LTP induction (Fig. 4H).

These data suggest that, in the barrel cortex, RWS triggers synaptic glutamate discharges that reach the bulk of local synapses. This appears consistent with the observation in acute slices that even mild activation of afferent fibres generates local iGluSnFR signals with no detectable release failures (*18*) that are otherwise prominent when firing is limited to just one presynaptic cell (*17*) (Fig. 1A-D). In fact, the *ΔF/F*_0_ iGluSnFR landscapes obtained here (Fig. 4C, fig. S4A) display virtually no ‘negative’ signal, suggesting a contiguous RWS-induced glutamate rise over at least 2-3 μm wide areas. Whether this occurs due to activation of TC synapses only or whether it also involves raised activity in local circuitry remains to be established.

Because the evoked *ΔF/F*_0_ iGluSnFR signal reflects glutamate that binds rapidly to the indicator, assessing receptor actions of free-diffusing glutamate is not straightforward. Firstly, astroglia-expressed iGluSnFR will compete for glutamate with high-affinity glial glutamate transporters, mainly GLT-1 type (*53, 70*). Because the on-rates of glutamate binding to iGluSnFR and GLT-1 are comparable (*71, 72*), and because both iGluSnFR and GLT-1 are co-expressed by astroglia, glutamate will bind to either in proportion to their local concentrations (cell-surface densities). Thus, the low local iGluSnFR/GLT-1 expression ratio could prevent detection of glutamate release by iGluSnFR, which is a possible reason why we found multiple TC synapses showing Ca^2+^ activity but no glutamate signal. By the same token, a dense patch of high-affinity receptors (such as NMDA or mGluR type) could be activated by glutamate at distances shown by the iGluSnFR detection. The prevalence and role of such volume-transmitted glutamatergic signals remain an open question. A significant reduction of local glutamate uptake capacity, either by withdrawal of perisynaptic astrocytic process (*40*) or localised activity-dependent inhibition of GLT-1 (*73*), could clearly boost such phenomena and a significant reduction of local glutamate uptake capacity could clearly boost such phenomena (*40*), which might play an important role in regulating cognitive functions of the human brain (*74, 75*).

## METHODS

### Organotypic Slice Preparation

Organotypic hippocampal slice cultures were prepared and grown with modifications to the interface culture method (*76*) from P6–8 Sprague-Dawley rats, in accordance with the European Commission Directive (86/609/EEC) and the United Kingdom Home Office (Scientific Procedures) Act (1986). Three hundred μm thick, isolated hippocampal brain slices were sectioned using a Leica VT1200S vibrotome in ice-cold sterile slicing solution consisting (in mM) of Sucrose 105, NaCl 50, KCl 2.5, NaH2PO4 1.25, MgCl2 7, CaCl2 0.5, Ascorbic acid 1.3, Sodium pyruvate 3, NaHCO3 26 and Glucose 10. Following washes in culture media consisting of 50% Minimal Essential Media, 25% Horse Serum, 25% Hanks Balanced Salt solution, 0.5% L-Glutamine, 28mM Glucose and the antibiotics penicillin (100U/ml) and streptomycin (100μg/ml), three to four slices were transferred onto each 0.4μm pore membrane insert (Millicell-CM, Millipore, UK), kept at 37ºC in 5% CO2 and fed by medium exchange for a maximum of 21 days in vitro (DIV).

### Biolistic Transfection of iGluSnFR variants

The second generation iGluSnFR variant SF-iGluSnFR.A184V, kindly gifted by Prof. Loren Looger was expressed in CA3 pyramidal cells in organotypic slice cultures using biolistic transfection techniques adapted from manafacturer’s instructions. In brief, 6.25 mg of 1.6 μm Gold microcarriers were coated with 30 μg of SF-iGluSnFR.A184V plasmid. Organotypic slice cultures at 5DIV were treated with culture media containing 5μM Ara-C overnight to reduce glial reaction following transfection. The next day cultures were shot using the Helios gene-gun system (Bio-Rad) at 120psi. The slices were then returned to standard culture media the next day and remained for 5-10 days before experiments were carried out.

### Axon tracing and 2PE imaging in axonal boutons

We used a Femtonics Femto2D-FLIM or a Femto3D-RC imaging system, integrated with patch-clamp electrophysiology (Femtonics, Budapest) and linked on the same light path to two femtosecond pulse lasers MaiTai (SpectraPhysics-Newport) with independent shutter and intensity control. Patch pipettes were prepared with thin walled borosilicate glass capillaries (GC150-TF, Harvard apparatus) with open tip resistances 2.5-3.5 MOhm. For CA3 pyramidal cells and dentate granule cells, the internal solution contained (in mM) 135 potassium methanesulfonate, 10 HEPES, 10 di-Tris-Phosphocreatine, 4 MgCl_2_, 4 Na_2_-ATP, 0.4 Na-GTP (pH adjusted to 7.2 using KOH, osmolarity 290–295), and supplemented with Cal-590 (300 μM; AAT Bioquest) for FLIM imaging in CA3 pyramidal cell axons.

Presynaptic imaging at CA3-CA1 synapses was carried out using an adaptation of presynaptic glutamate and Ca^2+^ imaging methods previously described (*17*). CA3 pyramidal cells were first identified as iGluSnFR expressing using 2PE imaging at 910 nm and patched in whole cell mode as above. Following break-in, 30-45 minutes were allowed for Cal-590 to equilibrate across the axonal arbour. Axons, identified by their smooth morphology and often torturous trajectory were followed in frame scan mode to their targets and discrete boutons were identified by criteria previously demonstrated to reliably match synaptophysin labelled punctae (*77*). In some cases, 4μM Alexa 594 was included with Cal-590 in the internal solution. The distinct two-photon excitation profiles of these two red emitting dyes enable morphology to be traced by Alexa 594 emission with 800nm excitation (Fig. 1A-B), and Ca^2+^ signals to be recorded in the same structure by Cal-590 emission at 910nm excitation (Fig. 1C), with no significant contribution of Alexa fluorescence to the Cal-590 emission.

To image glutamate release from mossy fibre giant boutons, dentate gyrus granule cells expressing iGluSnFR (using biolistic transfection, as described above) were patched, the axons were followed until putative giant boutons (main input to CA3 pyramidal cells) were identified as varicosities on axon collaterals with a width of at least 2-3 μm and one or more visible filopodial protrusions (*78, 79*). In separate experiments, granule cells were dialysed with the Ca^2+^ indicator OGB-1 (200 μM).

For fast imaging of action-potential evoked iGluSnFR, Cal-590, or OGB-1 fluorescence transients at single boutons, a spiral shaped (“Tornado”) scan line was placed over the bouton of interest (described further in the text) which was then scanned at a sampling frequency of ∼500 Hz with excitation at 910 nm. For multi-bouton imaging point-scans were made with a temporal resolution ∼333 or 250 Hz, usage described further in the text. Following a baseline period, action potentials initiated by brief positive voltage steps in voltage clamp mode (V_m_ holding -70 mV) were given with an interval of 50 ms.

### Cal-590 FLIM readout of Ca^2+^ concentration

Using the scanning methodologies described above line-scan data were recorded by both standard analogue integration in Femtonics MES and in TCSPC in Becker and Hickl SPCM using dual HPM-100 hybrid detectors. FLIM line scan data were collected as previously described (*17, 68*) and stored as 5D-tensors (t,x,y,z,T) to be analysed with custom written data analysis software available at (https://github.com/zhengkaiyu/FIMAS). Since morphological data was not of value when scanning only within the bouton x,y and z data were summed along their respective axes. For measurements of single trial baseline and action potential [Ca^2+^] data were then summed along the time axis with bin sizes adequate for accurate the use of the NTC measure to extrapolate [Ca^2+^] from FLIM data. multiple trials were summed to produce photon counts high enough to accurately determine dynamics of single action potential [Ca^2+^] transients.

### Imaging Uncaging evoked glutamate transients at single pre-synaptic boutons

Uncaging evoked glutamate transients were recorded in organotypic slices with bath application of 1.5 mM DNI-Glutamate on the Femto3D-RC imaging system (both Femtonics, Budapest) using protocols as described above. Spot uncaging was accomplished using the single galvo scanner pair by rapid switching from the spiral scan imaging modality to a point scan located off the edge of the pre-synaptic bouton of interest for 0.2 ms. During this time an uncaging pulse (λ_x_^2P^ = 720 nm, 10-18 mW depending on the depth of the bouton) was initiated by a 200 μs square wave voltage applied to a Pockels cell (Conoptics M302RM, Danbury, CT) in the light path of the uncaging laser. Once an uncaging-evoked glutamate transient was evoked, the duration and power of the pulse were calibrated with the aim to minimise the pulse width and best match the decay kinetics of action potential evoked glutamate transients whilst producing reliable responses (range of duration used 10-50 μs). The same protocol was then repeated four times with an interval of 100ms, for between 12 and 25 trials to match that using for measuring action potential evoked synaptic release.

### Dual transduction in the barrel cortex

All animal procedures were conducted in accordance with the European Commission Directive (86/609/EEC) and the United Kingdom Home Office (Scientific Procedures) Act (1986). Adult C57BL/6J mice, male and female, were transfected with two viral constructs, encoding Syn.jRGECO1a and GFAP.SF-iGluSnFR.A184S respectively. Mice were anaesthetised (isoflurane, maintenance at 1.5 - 2%), prepared for aseptic surgery and secured in a stereotaxic frame. Upon confirmation of the absence of pedal withdrawal reflex, two craniotomies of approximately 0.4 mm diameter were performed over the right hemisphere using a high-speed hand drill (Proxxon, Föhren, Germany), at sites overlying the posterior medial nucleus of the thalamus (POm) and the barrel cortex (S1BF). The entire microinjection into the POm was completed prior to performing the second craniotomy over S1BF. Stereotactic coordinates for POm injections (jRECO1a) were -2.2 mm and 1.2 mm along the anteroposterior and mediolateral axes, respectively. Two injection boluses were delivered at 2.9 and 3.1 mm beneath the dural surface. For S1BF injections (iGluSnFR), the bolus coordinates were -0.5 mm and 3.0 mm along the anteroposterior and mediolateral axes, respectively, delivered at a depth of 0.6 mm along an oblique approach angle to avoid tissue scarring in S1BF that might occlude the optical window. A warmed saline solution was applied to exposed cortical surface during the procedure.

Pressure injections of AAV9 hSyn.jRGECO1a (totalling 0.5 × 10^10^ genomic copies in a volume not exceeding 200 nL, initially supplied by Penn Vector Core, PA, USA; and later by Addgene, MA, USA) and AAV2/5.GFAP.iGluSnFR.A184S (0.1 × 10^10^ genomic copies, in a volume not exceeding 200 nL, kindly gifted by Prof. Loren Looger) were carried out using a glass micropipette at a rate of 1 nL sec-1, stereotactically guided to the POm and S1BF, respectively, as outlined above. Once delivery was completed, pipettes were left in place for 5 minutes before being retracted. The surgical wound was closed and the animal recovered in a heated chamber. Meloxicam (subcutaneous, 1 mg kg^-1^) was administered once daily for up to two days following surgery. Mice were subsequently prepared for cranial window implantation approximately 2 weeks later.

### Cranial window implantation

Mice were anaesthetised, prepared for aseptic surgery and secured in a stereotaxic frame as before during the viral transduction procedure. A large portion of the scalp was removed to expose the right frontal and parietal bones of the skull, as well as the medial aspects of the left frontal and parietal bones. The right temporalis muscles were reflected laterally to expose the squamous suture, to facilitate cement bonding during fixation of the cranial window implant. The exposed skull was coated with Vetbond (3M, MN, USA) and a custom-made headplate was affixed over the S1BF. The assembly was then secured with dental cement (SuperBond, Sun Medical Co. Ltd., Japan). Once the bonding agents had cured, the animal was removed from the stereotaxic frame and it’s headplate was secured in a custom-built head fixation frame. A craniotomy of approximately 4 mm diameter was carried out over the right somatosensory cortex, centred over the S1BF injection site. Immediately prior to removal of the skull flap, the surface was superfused with warmed aCSF (in mM; 125 NaCl, 2.5 KCl, 26 NaHCO3, 1.25 Na2HPO4,18 Glucose, 2 CaCl2, 2 MgSO4; saturated with 95% O2 / 5% CO2, pH 7.4). The dura was resected using a combination of 26G needles (pushed against a hard surface to introduce a curved profile), fine-tipped forceps (11252-40, Fine Science Tools, Germany) and 2.5 mm spring scissors (15000-08, Fine Science Tools, Germany), taking care not to penetrate to the pia mater. Once the dura was removed, a previously-prepared coverslip consisting of a 34 mm diameter round coverglass affixed beneath a 4 mm diameter round coverglass (Harvard Apparatus UK, affixed using a UV-curable optical adhesive (NOA61), ThorLabs Inc., NJ, USA) was placed over the exposed cortex. Slight downward pressure was applied to the coverslip using a stereotactically guided wooden spatula that was previously severed and sanded to allow some flexibility and preclude excessive force, as previously described (*80*). The superfusion was discontinued and excess aCSF was removed using a sterile surgical sponge, taking care not to wick fluid from beneath the cranial window. The coverslip was then secured with VetBond and dental cement, sequentially. Once cured, the animal was recovered in a heated chamber and returned to its homecage when ambulatory. Post-operative care was administered as before during the viral transduction procedure. Animals were allowed habituate to headplate implantation within their homecage for at least one week.

### Multiplexed 2PE imaging in vivo

Two-photon excitation was carried out using a wavelength multiplexing suite consisting of a Newport-Spectraphysics Ti:sapphire MaiTai tunable IR laser pulsing at 80 MHz and a Newport-Spectraphysics HighQ-2 fixed-wavelength IR laser pulsing at 63 MHz, as detailed earlier (*36, 68, 81*). The laser lightpaths were aligned (though not synchronised) before being point-scanned using an Olympus FV1000 with XLPlan N 25x water immersion objective (NA 1.05). Imaging was performed with head-fixed, awake animals as well as under lightly anaesthesia (fentanyl, 0.03 mg kg^-1^, midazolam, 3 mg kg^-1^, and medetomidine, 0.3 mg kg^-1^), with animals secured under the objective on a custom-built stage via the previously affixed headplate. First, exploratory acquisitions were performed with both lasers illuminating the tissue at 920 nm and 1045 nm, respectively, in order to locate thalamocortical axons in S1BF ramifying within the arbor of iGluSnFR-positive cortical astrocytes. Brief pulses of nitrogen were directed at the contralateral whiskers to confirm tactile responses within the regions of interest (rhythmic whisker stimulation, RWS; 5 seconds, 3 Hz). Measurements were performed throughout in L1 and L2/3, at depths of 50 - 150 nm. For high resolution axonal bouton sampling, an initial framescan of 4 - 20 Hz was performed, with a pixel dwell time of 2 μs and a mean laser power not greater than 30 mW at the focal plane. At this stage, tactile responses to short sensory stimulus trains (20 Hz, 200 milliseconds) were monitored at axonal boutons, both before and after potentiation while in Tornado scanning mode (400 – 500 Hz). Sensory-evoked synaptic potentiation (RWS-LTP) within the barrel cortex was induced as previously described (*28, 40*), via a sustained contralateral rhythmic whisker stimulation (120 sec, 3 Hz).

### Monte Carlo simulations of Brownian diffusion and cell surface binding

Monte Carlo algorithms for particle diffusion were designed and run with MATLAB: there were previously described in detail, and tested and constrained using various experimental settings (*62, 64, 82*). The simulation arena was a 4 μm wide cube, with 1000 particles ‘released’ instantaneously at the centre. For each value of free parameter Ψ (see below), 10 simulation trials were run, with different sphere distribution (generated so that *β* = 0.8 ± 0.025 over the trials). Particles positioned at time *t* at point **r**_*i*_ (*x, y, z*) were moved, over time step Δ*t*, to point **r**_*i*+1_(*x* + 2*δ*_*x*_Δ_1D_, *y* + 2*δ*_*y*_Δ_1D_, *z* + 2*δ*_*z*_Δ_1D_) where Δ_1D_ stands for the mean square displacement in the Einstein’s diffusion equation for 1D Brownian motion 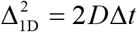, *D* = 0.65 μm^2^/ms is the glutamate diffusion coefficient in the interstitial space (*83*), and *δ*_*x*|*y*|*z*_ denotes a ‘delta-correlated’ (independently seeded, uncorrelated) uniform random number from the (−1, 1) range. The latter ensures that Brownian particles are equally likely to move into either direction whereas scale factor 2 for *δ* gives the average elementary displacement in *x-y-z* either −Δ_1D_ or +Δ_1D_. This algorithm provided the duty-cycle translational particle movements in a contiguous 3D space, over all directions with varied 3D steps, rather than over the rectangular 3D-lattice vertices used by us and many others previously. The randomness of the displacement vector helped avoid occasional numerical deadlocks for particles trapped near the space dead-ends formed by aggregated overlapped spheres. The time step Δ*t* (usually < 0.1 μs) was set to be small enough to prevent particles from ‘tunnelling’ through the smallest, 50 nm wide obstacles, and the actual value of *D* was verified at regular intervals.

The interaction with spheroids that represented either neuronal or astroglial fragments fell into the two corresponding cases. In the case of ‘neuronal’ spheroids, the interaction with diffusing glutamate particles was simulated as an elastic collision. In the case of ‘astroglial’ spheroids (which surface was populated with high-affinity transporters and/or iGuSnFR molecules), interaction was simulated as binding that occurred with the probability *P*. For each particle, *P* was a function of time *t* lapsed from its first collision with the current ‘astroglial’ spheroid, in accord with the basic lifetime expression for first-order reactions, *P* = 1− exp (*t* Ψ^−1^) where Ψ is the time constant (free parameter) that determines how likely is the oncoming binding event. Parameter Ψ thus combined, as a single quantity, the effects of binding affinity, binding-site surface density, and proximity to the diffusate. In practice, for each particle, computing *P* continued as long as the particle was within 5 nm of the ‘astroglial’ spheroid; it was reset to zero when it was further away (increasing the cut-off distance above 5 nm had negligible effect on *P*). Once bound, the particle remained bound to astroglial spheroids for the simulated time period (4-5 ms) because the characteristic time of glutamate unbinding for either GLT-1 or iGluSnFR is much longer (tens of milliseconds).

### Simulating sphere-filled space representing barrel cortex neuropil

There were at least two reasons to believe that randomly sized overlapping spheres would be a more realistic representation of neuropil compared to regular lattices of regular shapes, a tissue model used extensively by us and others previously. Firstly, multiple intersecting spheres give randomly shaped and randomly sized cellular elements and extracellular channels, as opposed to uniform or regular structures. Secondly, this approach provides a mixture of concave and convex shapes, including ‘diffusion dead-ends’ which are considered an important trait of brain neuropil (*84*). These realistic features of brain neuropil are not present in tissue models based on regular lattices.

Filling the space with the overlapping spheres followed the routines described in detail previously (*64*). In brief, the key parameter controlling this procedure was the volume fraction *β* occupied by the spheres: *β* = 1−*α* where *α* commonly stands for medium porosity, such as the volume fraction of the extracellular space in brain tissue. The *β* value was calculated by (a) scattering 10^5^ test points uniformly randomly throughout the arena, and (b) calculating the proportion of the point falling outside the spheres. We verified that increasing the number of such test points to 10^6^ altered *β* by <1%, pointing to asymptotic accuracy.

To fill the space with overlapping spheres that have a distributed size, we generated random 3D co-ordinates of sphere centroids across the simulation arena, and the random radius value for each sphere. The latter followed a uniform distribution between 50 and 300 nm, which roughly represented characteristic widths of cortical neuropil elements (fragments of neuronal and astroglial processes) seen in electron micrographs. For any new computation, the initial number of spheres was estimated and distributed based on their average volume and the average size (to give the required *β=0*.*8* value), and we left the co-ordinate origin unoccupied by any sphere. The space-filling cycle was repeated, with adjusted sphere numbers, until *β* approached the required value with ∼5% accuracy.

### Computing environment

Monte Carlo simulations were run using three computing environments. Firstly, a dedicated 8-node BEOWULF-style diskless PC cluster running under the Gentoo LINUX operating system (kernel 4.12.12), an upgraded version of that described earlier (*85*). Individual nodes comprised an HP ProLiant DL120 G6 Server containing a quad-core Intel Xeon X3430 processor and 8GB of DDR3 RAM. Nodes were connected through a NetGear Gigabit Ethernet switch to a master computer that distributes programs and collects the results on its hard disk. Secondly, on a UCL Myriad cluster: processors for each node, Intel(R) Xeon(R) Gold 6240 CPU @ 2.60GHz; cores per node 36 + 4 A100 GPUs; RAM per node 192GB, tmpfs 1500G, total 6 nodes. Thirdly, cloud computing with Amazon AWS: t4g.medium, memory 4GB. Parallelisation and optimisation of the algorithms and program codes were implemented by AMC Bridge LLC (Waltham, MA).

