## Supplementary Figure for "Pliable synaptic fidelity and excitatory inter-synaptic crosstalk in the intact brain"

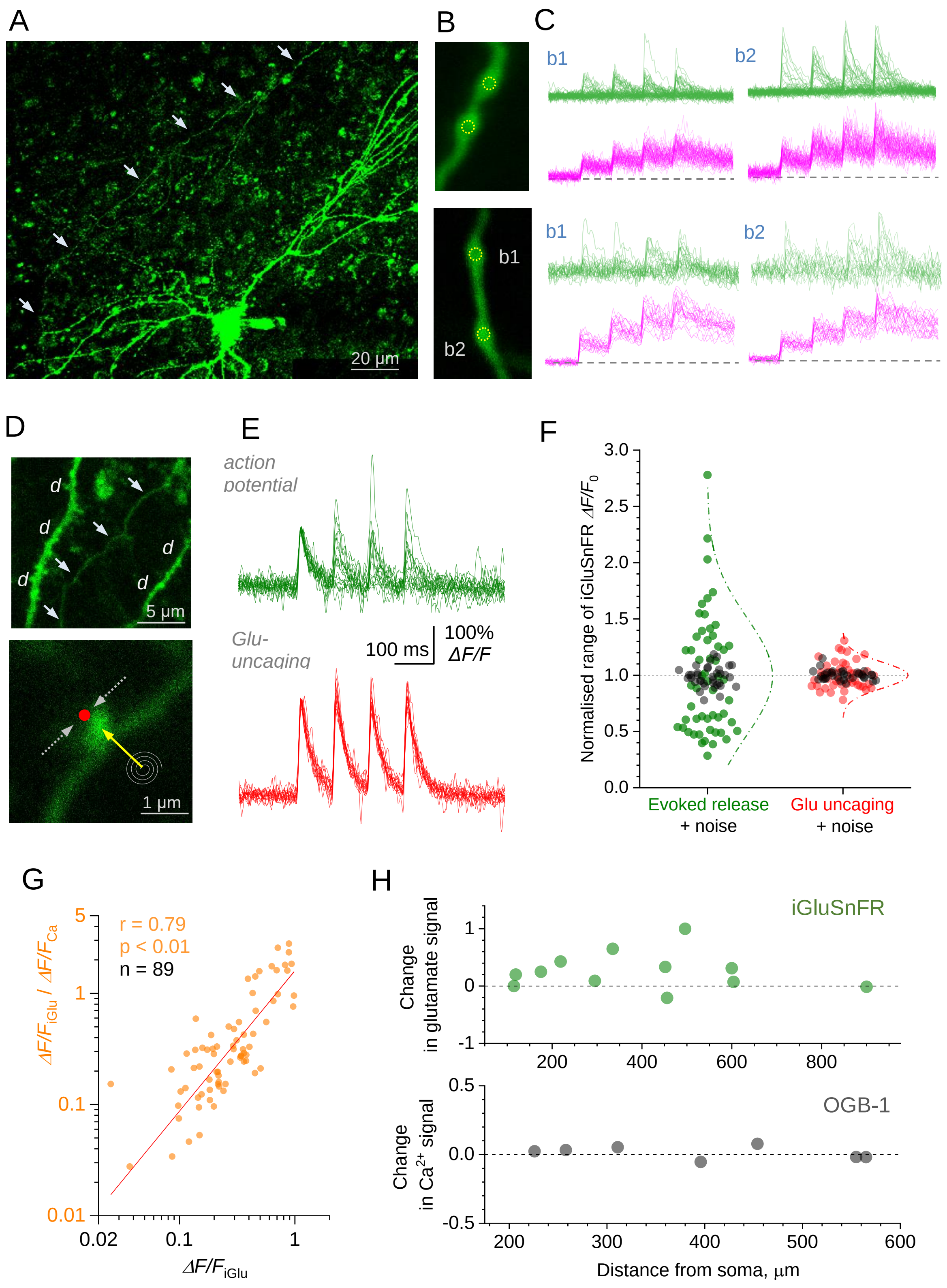


**Supplementary figure S1.** Controls for the Glu/Ca ratio as a release probability tracker.

(A) CA3 pyramidal cell (organotypic hippocampal slice) expressing SF-iGluSnFR.A184V and dialysed whole-cell with Cal-590, with 4 ROIs along the axon (arrowheads) traced in the iGluSnFR channel (collage of 10-15 μm deep image stack projections, $\lambda_x^{2P}$=910); scattered iGluSnFR expression and autofluorescence somewhat contaminates the image.

(B) Traced presynaptic axonal boutons, with multi-point scan positions as illustrated.

(C) Fluorescence time course (integrated point-scan signal at 500 Hz) in iGluSnFR (green) and Cal-590 (magenta) channels ($\lambda_x^{2P}$=910 nm), in response to four action potentials 50 ms apart, as indicated, for the corresponding boutons shown in B; 20-25 trials shown overlapped.

(D) Control experiment comparing iGluSnFR responses to synaptic discharges of glutamate versus a glutamate spot-uncaging pulses; top, fragment of an iGluSnFR-expressing CA3 pyramidal cell, with fragments of dendrites (*d*) and the axon (arrowheads); bottom, axonal bouton of interest, showing positioning of the Tornado scan (spiral, arrow) and spot-uncaging (red dot, dotted arrows).

(E) Example of iGluSnFR responses to four action potentials (top, green; normalised to 1st successful response; traces with all-failure responses were ignored) and glutamate spot-uncaging (red, right) 100 ms apart, in the experiment shown in D.

(F) Normalised (to one) amplitude distributions of $\Delta F/F$ iGluSnFR responses to action potential (left, green dots; release failures ignored) and glutamate spot-uncaging (red dots, right), with baseline noise (black dots); dash-dotted lines, approximating normal distributions indicating standard deviation of 1.18 (evoked response) and 0.36 (spot-uncaging).

(G) The ratio of $\Delta F/F$ iGluSnFR response (250 ms over four action potentials) versus $\Delta F/F$ Cal-590 response ('Glu/Ca' ratio, orange), plotted against $\Delta F/F$ iGluSnFR response (direct glutamate release readout, n = 89 boutons) at 2 mM extracellular [$Ca^{2+}$]; straight line, linear regression; r, coefficient of correlation; log-log scale.

(H) Mossy fibre experiments (Fig. 1G-I): somatic depolarisation induced changes in the $\Delta F/F$ iGluSnFR signal (green, top) and $\Delta F/F$ OGB-1 signal (grey, bottom), plotted against bouton distance from the soma.

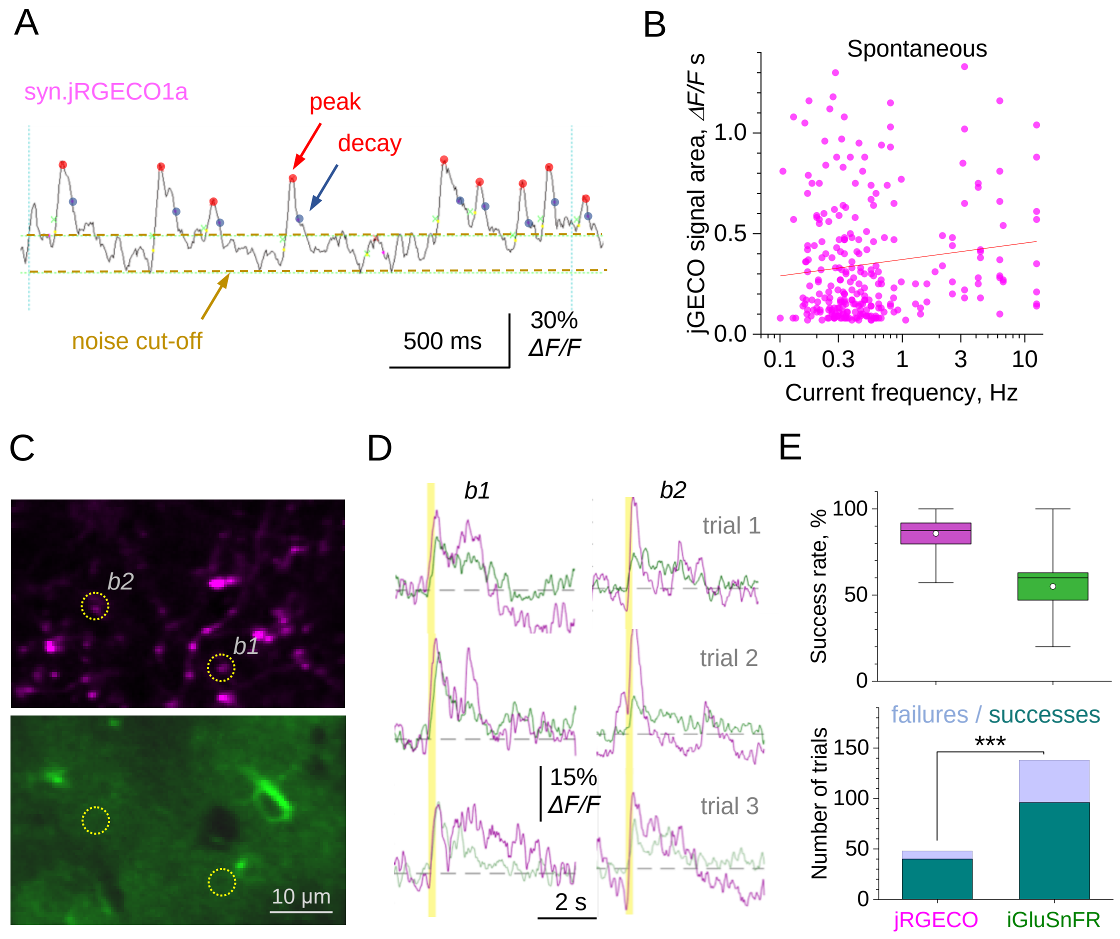


**Supplementary figure S2**. Spontaneous and evoked presynaptic $Ca^{2+}$ activity in thalamocortical axonal boutons.

(A) Example, analysis of spontaneous presynaptic $Ca^{2+}$ activity events (jRGECO1a channel; Mini Analysis 6, Synaptosoft). Main filtering parameters: baseline average window, 100 ms; peak search, 200 ms; decay time search, 500 ms; decay time point, $e^{-1}$; detection cut-off, ~2SD of baseline noise.

(B) Strength of spontaneous $Ca^{2+}$ activity events (area under the $\Delta F/F_0$ jRGECO1a curve) plotted against the ongoing event frequency (inverse timespan between the current and previous event, n = 263); line, linear regression (Pearson's correlation coefficient R = 0.13, p = 0.013).

(C) Multiplexed imaging of $Ca^{2+}$ activity (jRGECO1a-expressing axonal boutons, top) and glutamate release (iGluSnFR-expressing astroglia channel, bottom) in the barrel cortex; b1 and b1, two axonal boutons showing responses to whisker stimuli.

(D) Examples of fluorescence responses (magenta jRGECO1a, green iGluSnFR) recorded from boutons 1 and 2 in C, in response to 200 ms whisker stimuli (vertical segment), as indicated.

(E) Top: success rate (proportion of fluorescence responses detected above noise in all trials), in jRGECO1a (136 trials) and iGluSnFR channels (50 trials); dots, mean values; horizontal line, median; box, ± SE. Bottom: proportions of failures / successes in the two channels, as indicated; ***, $p = 0.00678$ (Proportion test, $Z=2.707$).

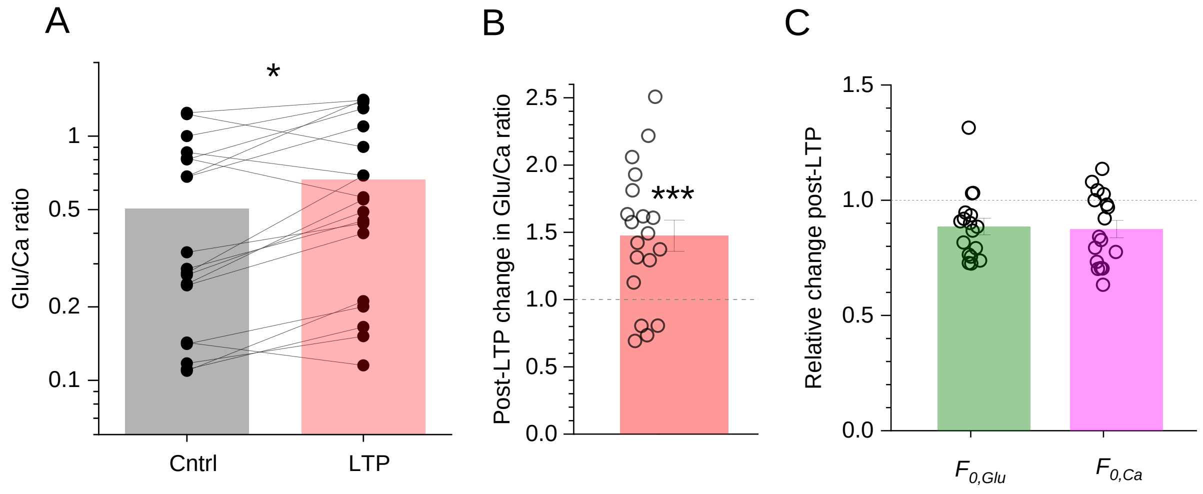


**Supplementary figure S3**. Rhythmic-whisker-stimulation induced long-term potentiation (RWS-LTP) of transmission in the barrel cortex increases fidelity while reducing excitation of individual thalamocortical connections.

(A) The Glu/Ca ratio (ratio between $\Delta F/F_0$ responses in the iGluSnFR and jRGECO channels, as shown in Fig. 3C) in control conditions (Cntrl) and 10-30 min after RWS-LTP induction (LTP); dots, individual bouton data (n = 19 from 6 animals); lines connect same-bouton data; bars, mean value; ordinate, log scale; *, p = 0.016 (paired-sample t-test), p = 0.020 (non-parametric paired-sample sign test); same dataset as in Fig. 3D.

(B) Relative change in the Glu/Ca ratio 10-30 min after RWS-LTP induction; dots, individual boutons (n = 19); bar, mean ± SEM; ***, p < 0.001; same dataset as in Fig. 3D.

(C) Relative change in the baseline fluorescence ($F_0$) at thalamocortical boutons in the iGluSnFR channel ($F_{0,Glu}$) and jRGECO1a channel ($F_{0,Ca}$); dots, individual boutons (n = 19 from 6 animals); bars, mean ± SEM.

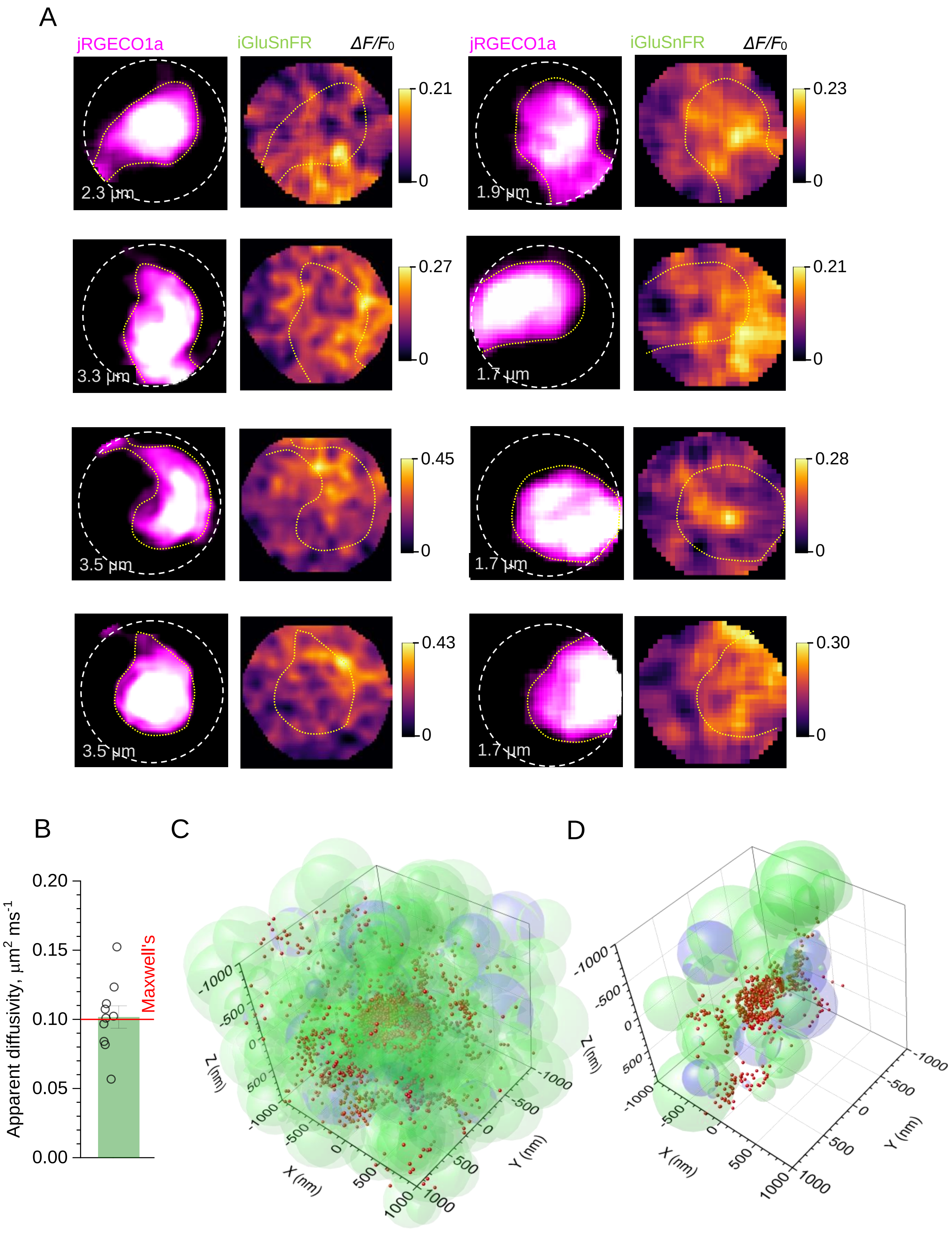


**Supplementary figure S4.** Extrasynaptic glutamate escape in thalamocortical (TC) synapses activated by whisker stimulation.

(A) Left panels (magenta): examples of high-resolution Tornado-linescan multiplexed imaging of TC axonal boutons, showing a jRGECO1a-expressing TC axonal bouton; dotted circles, Tornado scan area coverage (diameter shown); yellow dotted line, axonal bouton as judged by the fluorescence of cytosolic jRGECO1a. Right panels (false $\Delta F/F_0$ colour scale shown): the $\Delta F/F_0$ iGluSnFR signal landscapes, generated as the $\Delta F/F_0$ image ratio where $\Delta F$ and $F_0$ are the landscape images of iGluSnFR-expressing astroglia recorded during 200 ms before and 200 ms during a 200 ms long rhythmic whisker stimulation, as shown in Fig. 4A-B.

(B) Control tests for the model of glutamate release and free diffusion in a porous neuropil represented by a random scatter of unequal overlapping spheres: comparison of Monte Carlo simulation outcomes (dots, individual trials; bar, mean ± SEM) with the theoretical apparent diffusivity $D_{app}$ in a porous medium (horizontal line; Maxwell's equation $D_{app} = \frac{2\alpha D_{free}}{3-\alpha}$ where $\alpha$ is medium porosity). Main parameters: simulation arena, 4 µm wide cube; release site, centre; number of Brownian particles 1000; free diffusivity $D$ = 0.5 $\mu m^2$/ms; space fraction occupied by spheres, 80% ($\alpha$ = 0.2). Notations: dots, individual simulation trials; bar, average ± SEM.

(C) Simulation snapshot of release (centre of the arena), diffusion, and binding of glutamate molecules (red dots) to astroglia-expressed iGluSnFR (and glutamate transporters GLT-1), in a neuropil modelled *in silico* by a scatter of overlapping spheres (text and Methods); green spheroidal shapes, neuronal elements; blue spheroidal shapes, astroglial elements; all spheroidal shapes rendered semi-transparent for illustration; other parameters as in Fig. 4F.

(D) Simulation snapshot as in C, but with the graphic representation limited to a narrow slab (200 nm along the x-axis), for clarity purposes; other notations as in C.
